# Cervicovaginal Dysbiosis in HPV-Negative Women: Metagenomic Evidence Implicates *Achromobacter* in Female Infertility

**DOI:** 10.64898/2026.03.23.713732

**Authors:** Hazrat Ali, Shiful Islam Sujan, Kamrun Nahar, Firoz Ahmed, Nafisa Azmuda, Salma Akter, Nihad Adnan

## Abstract

The cervicovaginal microbiome is pivotal to reproductive health, yet its dynamics in HPV-negative women with gynaecological disorders remain underexplored. We investigated microbial diversity and taxonomic shifts in HPV-negative women from Bangladesh using 16S rRNA gene sequencing and shotgun metagenomics. Of 224 women screened, 136 were HPV-negative; 29 underwent 16S profiling, and three infertility-associated cases were further analyzed by shotgun metagenomics. Healthy controls exhibited low alpha diversity and a *Lactobacillus*-dominated profile (98.2%), reflecting ecological stability. In contrast, pathological cases displayed significantly elevated richness and evenness, reduced *Lactobacillus* (28.0%), and enrichment of anaerobic and opportunistic taxa, including *Bifidobacterium* (23.4%), *Achromobacter* (12.9%) and *Sneathia* (7.5%). Distinct microbial signatures emerged across clinical subgroups: pelvic inflammatory disease was enriched in *Bifidobacterium*, intra-menstrual bleeding retained moderate *Lactobacillus*, while infertility exhibited prominent dominance of *Achromobacter* (45.5%). Shotgun metagenomics confirmed *Achromobacter spp.* (*A. ruhlandii, A. dolens, A. xylosoxidans*) as the predominant taxa (84.9%) in infertility cases, accompanied by depletion of protective *Lactobacillus*. Functional inference revealed conserved metabolic backbones but disease-specific enrichment of stress-response and biosynthetic pathways, particularly in infertility and PID. Co-occurrence network analysis identified condition-specific microbial consortia, with *Achromobacter* forming infertility-associated clusters. This study represents the first integrated application of amplicon and shotgun metagenomic approaches to profile the cervicovaginal microbiota in HPV-negative women. It identifies *Achromobacter* as a potential microbial biomarker of infertility and highlights the urgent need for microbiome-informed diagnostics and targeted interventions to restore cervicovaginal homeostasis.

## 1. Introduction

The human vaginal ecosystem harbours one of the most diverse and lively microbial communities in the body, known as the vaginal microbiota (VMB). This community plays a pivotal role in maintaining reproductive health by producing lactic acid, hydrogen peroxide, and bacteriocins that inhibit pathogen colonization. Disruptions in VMB, called dysbiosis, have been linked to over 70% of gynaecological issues, including bacterial vaginosis (BV), pelvic inflammatory disease (PID), and infertility (1–3). Although HPV-related dysbiosis has been widely studied, the microbial changes in HPV-negative women are less understood, especially in low-resource areas like Bangladesh.

The normal vaginal microbiota comprises bacterial species that maintain homeostasis and prevent sexually transmitted infections (2–4), (5). Researchers have identified more than 50 species of microorganisms in the vaginal ecosystem (5). A typical balanced VMB is typically dominated by *Lactobacillus* species, which constitute 70–95% of the normal flora (3,5–7) and are classified into five Community State Types (CSTs), i.e., CSTs I (*L. crispatus*), II (*L. gasseri*), and V (*L. jensenii*) are considered protective, while CSTs III (*L. iners*) and IV (diverse anaerobes) are associated with dysbiosis (8,9). Beyond *Lactobacillus*, other taxa such as *Gardnerella vaginalis, Prevotella spp., Atopobium vaginae,* and *Sneathia spp.* are frequently implicated in vaginal disorders (10,11). The composition of the VMB fluctuates with age, hormonal status, sexual activity, contraceptive use, and ethnicity, which may be normal or pathological (12). Exogenous factors such as douching, lubricants, smoking, and intrauterine devices also modulate microbial diversity (10,13).

Studies have shown that, during the prepubertal stage, the VMB is dominated by *Escherichia coli*, diphtheroids, and coagulase-negative staphylococci. The vaginal tract of a premenopausal woman contains more than 20 species of *Lactobacillus*, while reproductive-age women exhibit VMB dominated by one or two *Lactobacillus* species, most commonly *L. crispatus*, *L. jensenii*, *L. gasseri*, and *L. iners* (2,5,14–16). The number of Lactobacilli decreases after menopause due to decreased estrogen levels (2,5,16,17). During menstruation, a shift is characterized by a 100-fold decrease in *L. crispatus* and an increase in *L. iners*, *G. vaginalis*, *Prevotella bivia*, and *Atopobium vaginae.* (2,5,16,17). Studies have also shown that oral contraceptives containing medroxyprogesterone, copper-IUDs, and douches reduce Lactobacillus colonization, whereas lubricants containing nonoxynol-9 (N-9) may increase E. coli colonization (5,18–20).

Any alteration or disruption in the normal VMB is termed dysbiosis, which is characterized by a decline in *Lactobacillus* and enrichment of anaerobes such as *Gardnerella*, *Prevotella*, *Atopobium*, and *Sneathia*. Dysbiosis has been linked to bacterial vaginosis (BV), pelvic inflammatory disease (PID), infertility, and increased susceptibility to sexually transmitted infections. Studies have shown that, under disease conditions, vaginal microbiomes are characterized by increased microbiome diversity and enriched colonization of anaerobic bacteria, like the enrichment of *Sneathia, Megasphaera, Prevotella, Gardnerella, Atopobium, Streptococcus,* and *Ureaplasma* (21,22), (2,23–25). BV alone is three times more prevalent in infertile women and is associated with adverse reproductive outcomes, including miscarriage and preterm birth (26,27), (28,29). PID, often caused by *Chlamydia trachomatis* and *Neisseria gonorrhoeae*, can lead to ectopic pregnancy and long-term infertility (30,31).

While the role of human papillomavirus (HPV), particularly high-risk genotypes like HPV-16 and HPV-18, in driving cervicovaginal dysbiosis and cancer progression is well documented, microbial dynamics in HPV-negative women remain poorly defined (32). This gap is especially critical in low-and middle-income countries such as Bangladesh, where the burden of gynaecological disorders is high but microbiome-based diagnostics are limited.

To address this, we characterized the cervicovaginal microbiota of HPV-negative women presenting with diverse gynaecological conditions using both 16S rRNA gene sequencing and shotgun metagenomics. By linking microbial diversity and community composition to specific pathological conditions, this study identifies microbial signatures of dysbiosis. It highlights potential biomarkers, such as *Achromobacter*, with implications for early diagnosis and microbiome-targeted interventions.

## 2. Materials and Methods

### 2.1. Study Population and Sample Collection

This prospective study was conducted between 2022 and 2023 at the Virology and Histopathology Department of Memorial KPJ Specialized Hospital, Dhaka, Bangladesh, in collaboration with the Department of Microbiology at Jahangirnagar University, Bangladesh. A total of 224 women aged 21–51 years presenting to the gynaecology outpatient clinic for HPV screening or Papanicolaou (Pap) testing were enrolled following written informed consent. Cervicovaginal swab specimens were collected using sterile cytobrushes, inserted approximately 0.5 cm into the endocervical canal, rotated counterclockwise three times, and then applied to the ectocervical surface. To minimize contamination, contact with the external genitalia and vaginal introitus was avoided. Samples were transferred to −80°C for preservation before further characterization.

### 2.2. HPV Screening and Cytological Evaluation

All samples were screened for HPV using real-time qPCR with the GeneProof Human Papillomavirus Screening Kit. HPV-negative samples were further evaluated for cytological abnormalities via Pap testing. From these, 29 HPV-negative samples were randomly selected and stratified into two groups based on clinical and cytological findings: Control (n = 7) and Pathological (n = 22). The Pathological group was further categorized into five clinical subgroups: discharge, infertility, pelvic inflammatory disease (PID), intra-menstrual post-vaginal bleeding (IPVB), and ‘Others’. The Control group was designated as “Control” for comparative analysis.

### 2.3. DNA Extraction

Genomic DNA was extracted from the 29 selected samples using the QIAamp DNA Mini Kit (Qiagen, UK), following the manufacturer’s protocol. DNA purity and concentration were assessed using a NanoDrop spectrophotometer, with A260/A280 ratios used to confirm sample integrity.

### 2.4. 16S rRNA Gene Amplicon Sequencing

To profile bacterial community composition, 16S rRNA gene amplicon sequencing targeting the V3-V4 hypervariable regions was performed by EzBiome (Gaithersburg, MD, USA). DNA concentration was quantified using a Qubit fluorometer (Thermo Fisher Scientific, USA). PCR amplification was conducted using primers 341F (5′-CCTACGGGNGGCWGCAG-3′) and 806R (5′-GACTACHVGGGTATCTAATCC-3′) (33,34). Amplicons were sequenced using paired-end 2 × 300 bp chemistry on the Illumina MiSeq platform (35).

### 2.5. Bioinformatic and Statistical Analysis of 16S rRNA Gene Data

Raw sequencing reads were initially assessed for quality using FastQC (v0.12.1), and low-quality bases (Q < 20) together with adapter sequences were trimmed using Trimmomatic (v0.39) (36), (37). Subsequent processing was performed in QIIME2 (v2024.10) with VSEARCH integration, including paired-end joining, denoising, chimaera removal, and *de novo* clustering at 99% sequence identity (38), (39). Taxonomic assignment was conducted using a Naïve Bayes classifier trained on the Greengenes2 reference database (v2024.09) (40). Downstream microbial community analyses were performed in R (v4.5.2, RStudio v2026.01.0+392) using established packages: *phyloseq* (v1.52.0), *vegan* (v2.7.2), *DESeq2* (v1.48.2), *ggplot2* (v4.0.2), *ggpubr* (v0.6.2), *dplyr* (v1.2.0), *pheatmap* (v1.0.13), *corrplot* (v0.95), and *VennDiagram* (v1.8.2). Alpha diversity indices (Chao1, Shannon, Simpson, Fisher’s alpha, and Pielou’s evenness) were calculated from ASV tables rarefied to 16,794 reads, while statistical analyses were conducted on normalized, non-rarefied data. Normality and homogeneity of variance were assessed using Shapiro-Wilk and Levene’s tests, respectively. Depending on the data distribution, group comparisons used ANOVA or the Kruskal-Wallis test, with Dunn’s post hoc or the Wilcoxon rank-sum test for pairwise comparisons. All p-values were adjusted for multiple testing using the false discovery rate (FDR), and effect sizes for two-group comparisons were quantified via rank-biserial correlation (significance threshold: FDR-adjusted p < 0.05). Beta diversity was estimated using Bray-Curtis dissimilarity on relative abundance-transformed ASV tables. PERMANOVA (adonis2, 9999 permutations) was applied with stratification for repeated measures, and dispersion homogeneity was verified using betadisper. Multivariable models incorporated age and BMI as covariates. Pairwise PERMANOVA with FDR correction was used for multi-group comparisons. Ordination analyses included PCoA, NMDS, and DCA, with 95% confidence ellipses; compositional patterns were further validated using CLR-transformed Aitchison distances. A fixed random seed (set.seed(123)) ensured reproducibility across analyses. Taxonomic composition and core microbiome analyses were performed, and differential abundance testing was conducted using DESeq2, with significance defined as FDR-adjusted p < 0.05 and |log□ fold change| > 1. Functional potential of the cervicovaginal microbiome was inferred using PICRUSt2 (v2.6.2). Representative OTUs were normalized by 16S rRNA gene copy number, and metagenomic functional profiles were predicted against the MetaCyc pathway database. Predicted pathway abundances were compared across clinical groups, visualized via functional heatmaps, and integrated into downstream analyses. Taxon-taxon correlations were computed using Pearson correlation with FDR adjustment.

### 2.6. Shotgun metagenomic sequencing and taxonomic profiling

To comprehensively characterize the cervicovaginal microbiome, particularly in infertility-associated samples, shotgun metagenomic sequencing was performed on three representative infertility samples, selected from the 29 HPV-negative cases. Genomic DNA was processed and sequenced by EzBiome (Gaithersburg, MD, USA) on the Illumina MiSeq platform using proprietary protocols. Taxonomic profiling was conducted using Kraken2 with a pre-built k-mer database (k = 35) from EzBioCloud and NCBI RefSeq (41), (42). Reads were aligned to a custom Bowtie2 database using the very-sensitive option and Phred33 quality threshold (43). BAM files were processed with Samtools (44), and coverage metrics were calculated using Bedtools (45). Species-level abundance was quantified using an in-house script, considering taxa with ≥25% core gene coverage (bacteria and archaea) or genome coverage (fungi and viruses). Relative abundances were normalized to reference genome lengths.

## 3. Results

### 3.1 Study Population and Screening Outcomes

A total of 224 women aged 21-51 years were enrolled for HPV screening and cytological evaluation. Among them, 136 (60.7%) tested HPV-negative, while 88 (39.3%) were HPV-positive. HPV-18 emerged as the most prevalent high-risk genotype, followed by other high-risk variants, whereas HPV-16 was particularly absent in this cohort. From the HPV-negative cohort, 29 cervicovaginal samples were selected for microbiome profiling and stratified into a Control group (n = 7) and a Pathological group (n = 22). The pathological group included women presenting with vaginal discharge, infertility, pelvic inflammatory disease (PID), intra-menstrual post-vaginal bleeding (IPVB), and other gynaecological symptoms. Control participants were predominantly aged 26-35 years, whereas pathological cases spanned a broader age range and exhibited higher frequencies of overweight/obesity and comorbidities such as hypothyroidism, hypertension, and diabetes. Within the pathological group, infertility and bacterial vaginosis (BV) were the most common clinical diagnoses. These demographic and clinical variations provided a diverse framework for investigating cervicovaginal microbial dynamics across health and disease states (Figure 1).

**Figure 1.**
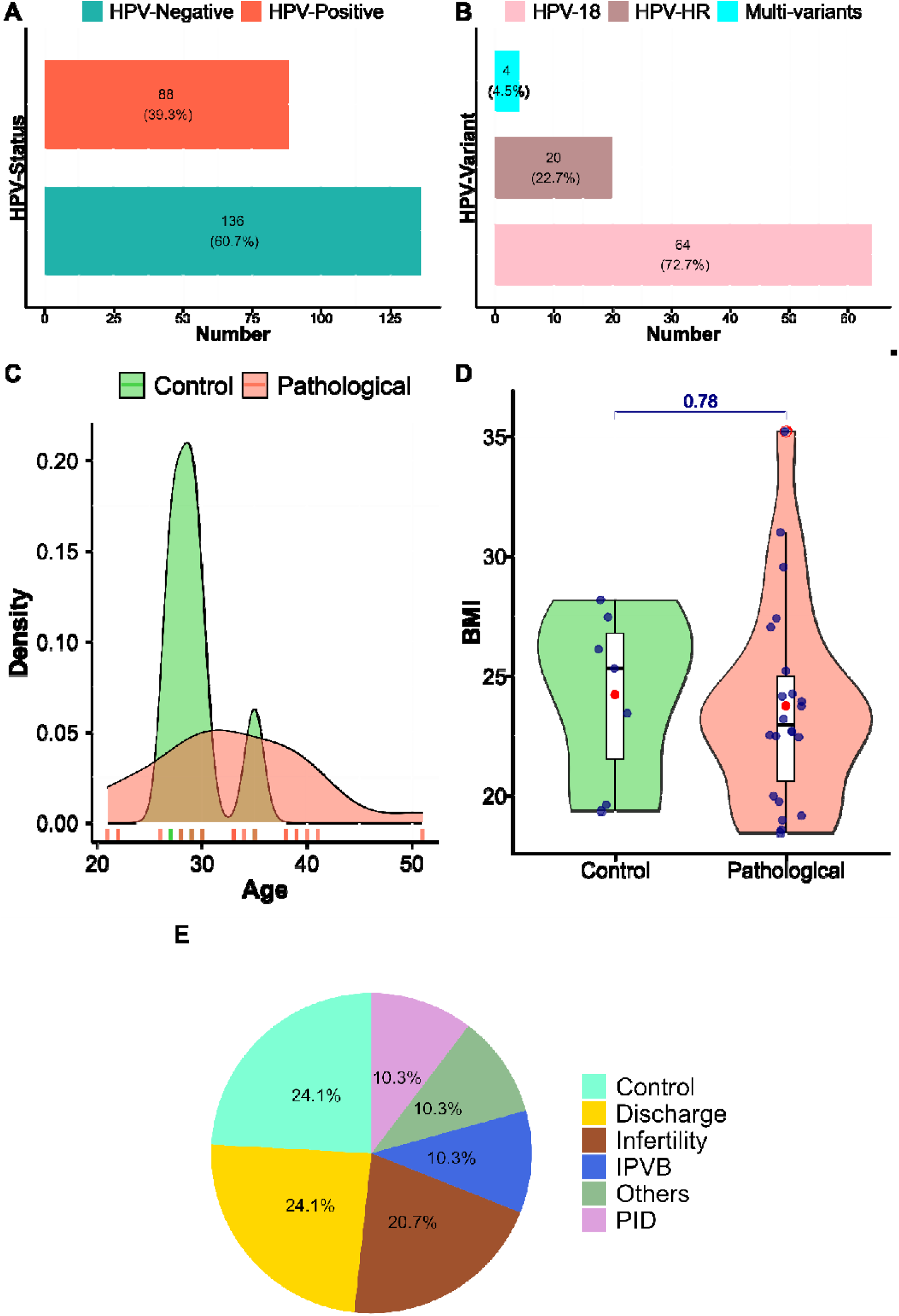
Clinical and demographic overview of the study population.

### 3.2 Diversity Patterns of the Cervicovaginal Microbiome

#### Alpha diversity and Beta diversity

Microbial richness and evenness differed significantly between healthy Controls and Pathological groups. Across multiple indices, including Shannon, Simpson, Fisher’s alpha, and Pielou’s evenness, pathological samples exhibited consistently higher alpha diversity than Controls. Shannon diversity was significantly elevated in pathological cases (p = 0.015), suggesting more complex, evenly distributed microbial communities. Similarly, Simpson and Pielou’s evenness indices demonstrated significant increases (p = 0.004 and p = 0.005, respectively), reinforcing the transition toward a heterogeneous microbial landscape. In contrast, Control samples displayed low diversity, consistent with a stable, *Lactobacillus*-dominated ecosystem typical of healthy cervicovaginal environments. Among pathological subgroups, infertility and PID cases contributed most strongly to increased diversity, whereas discharge and IPVB cases exhibited intermediate profiles, suggesting pathological conditions disrupt ecological stability, facilitating colonization by diverse anaerobic and opportunistic taxa (Figure 2A).

**Figure 2.**
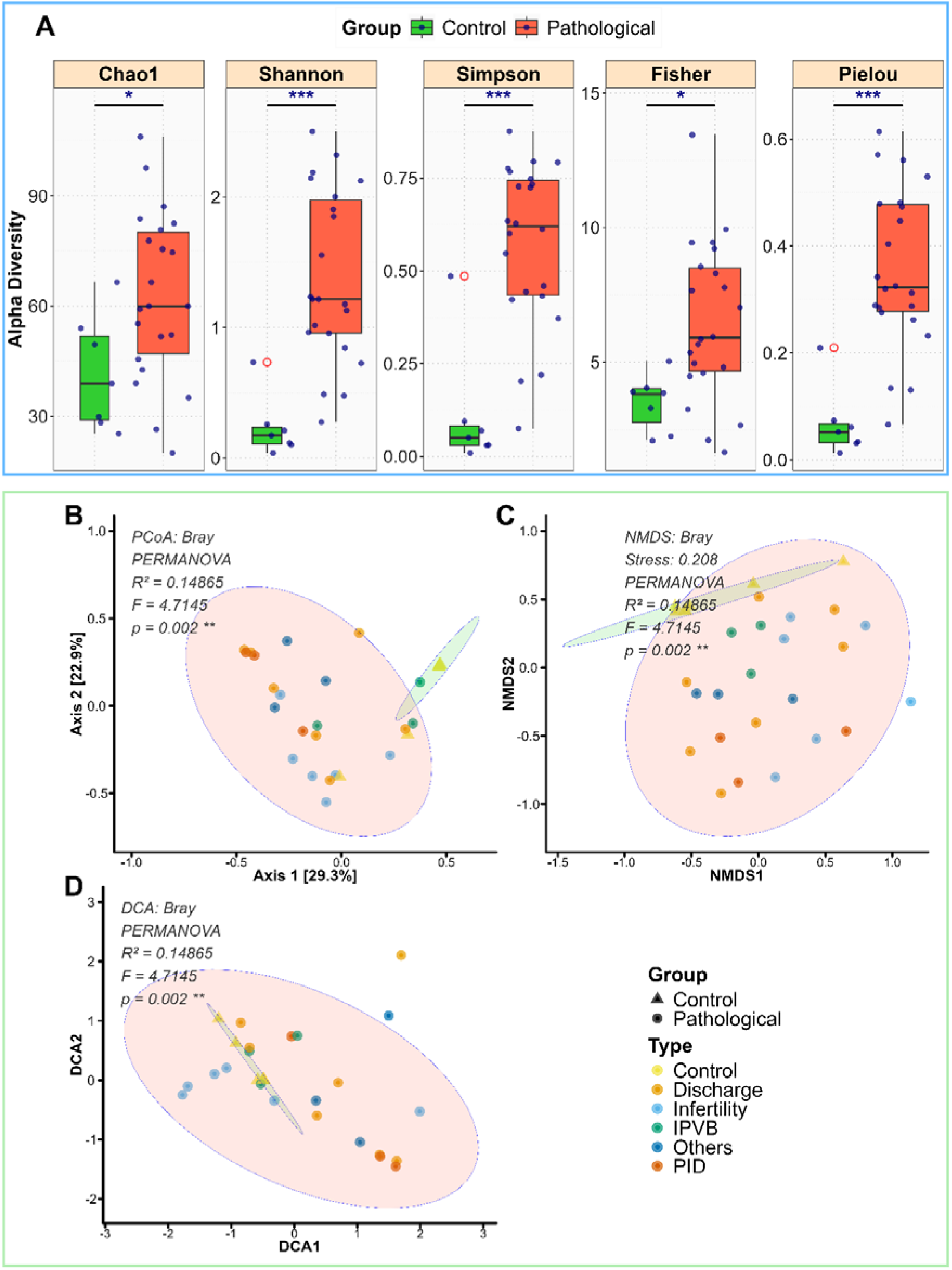
Alpha and Beta Diversity of Cervicovaginal Microbiomes Across Control and Pathological Groups.

Community composition also differed significantly between groups. Bray-Curtis dissimilarity-based PERMANOVA confirmed separation between Controls and Pathological groups (R² = 0.149, F = 4.716, p = 0.0005). Pairwise analysis supported distinct clustering of Pathological samples relative to Controls. Importantly, adjusting for age and BMI in multivariable PERMANOVA retained significant group effects (R² = 0.153, F = 4.939, p = 0.0003), while age and BMI themselves were not significant (p > 0.14). Group dispersion analyses further highlighted heterogeneity. betadisper ANOVA revealed significant differences in dispersion (F = 15.70, p = 0.00049; permutation test p = 0.001), with Pathological samples showing greater within-group variability (mean distance to centroid = 0.576) than Controls (0.240). CLR-transformed Aitchison distances confirmed compositional trends, though marginally non-significant (p = 0.093), supporting that observed differences were not solely compositional artifacts.

Ordination analyses aligned with these findings. PCoA of Bray-Curtis distances revealed clear clustering of Control and Pathological samples, with 95% confidence ellipses demonstrating separation. NMDS produced a stable two-dimensional solution (stress = 0.208), showing overlapping yet distinguishable patterns consistent with PCoA. DCA further highlighted gradients associated with pathological subtypes, with infertility and PID samples driving the largest deviations (Figure 2B–D).

### 3.3 Taxonomic Composition of the Cervicovaginal Microbiome and Shifts Across Clinical Conditions

#### 3.3.1 Phylum- and Class-Level Patterns

Analysis of the cervicovaginal microbiome revealed pronounced differences in taxonomic composition between healthy Controls and Pathological samples. At the phylum level, healthy Controls were overwhelmingly dominated by Bacillota (98.4%), with minor contributions from Pseudomonadota (1.4%) and other phyla (<0.1%). In contrast, Pathological samples displayed a clearly more diverse community, with Bacillota reduced to 36.2% and substantial representation of Pseudomonadota (26.3%) and Actinomycetota (26.3%), alongside elevated levels of Fusobacteriota (7.5%) and Bacteroidota (3.1%). These findings indicate a clear ecological shift from the Bacillota-dominated healthy state toward a heterogeneous microbiome in disease conditions (Figure 3).

**Figure 3.**
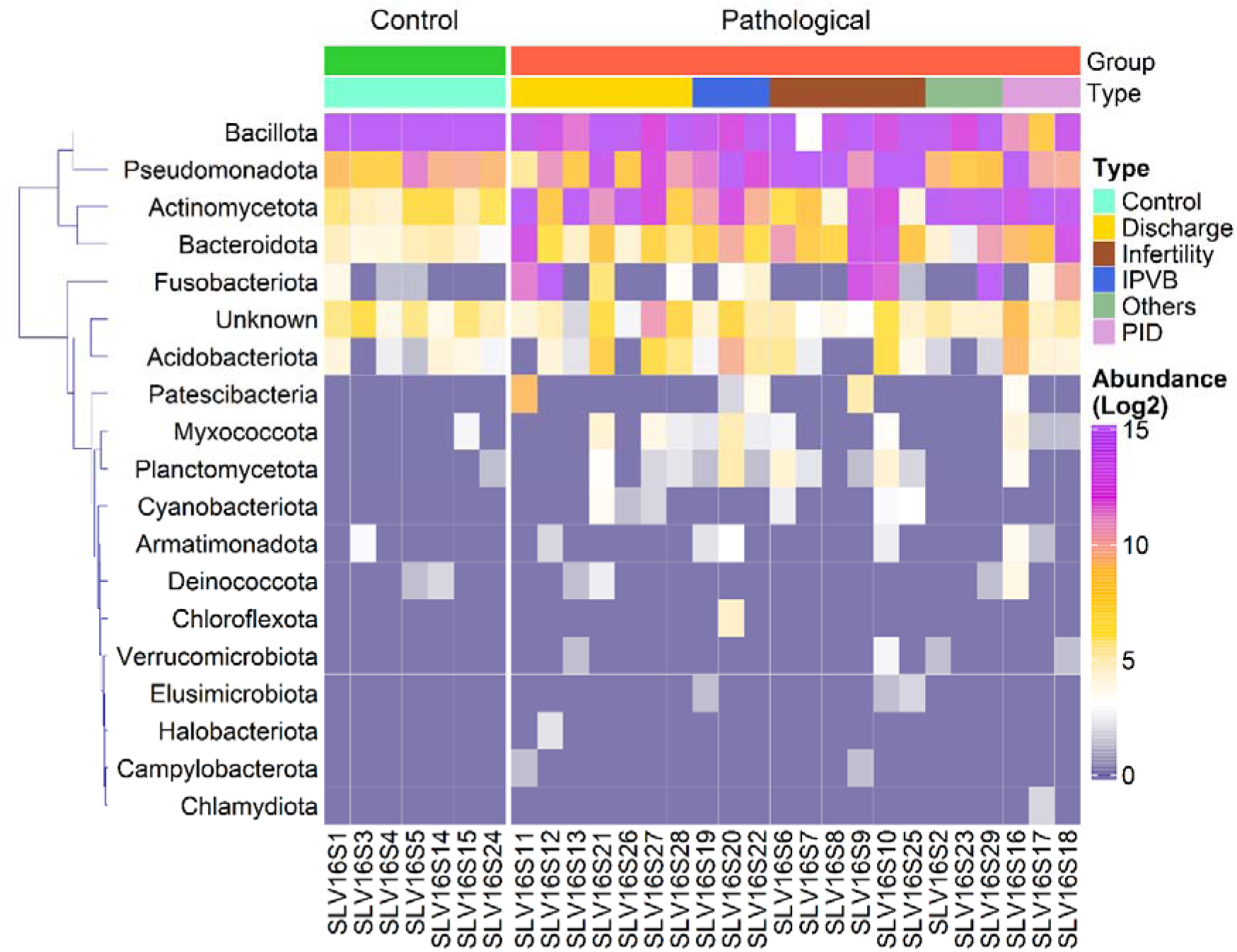
Phylum-Level Microbiome Composition Across Cervicovaginal Clinical Types.

At the class level, the cervicovaginal microbiome displayed distinct compositional shifts between healthy and pathological groups. Control samples were almost exclusively dominated by Bacilli (98.4%), with only minor contributions from Alphaproteobacteria (0.8%) and Gammaproteobacteria (0.6%), reflecting the stability of a *Lactobacillus*-driven ecosystem. In contrast, pathological samples exhibited markedly reduced Bacilli (30.4%) and notable enrichment of Actinomycetes (24.6%), Gammaproteobacteria (13.7%), Alphaproteobacteria (12.6%), and Fusobacteriia (7.5%), indicating a transition toward a heterogeneous and dysbiotic community structure. Stratification by clinical subtype further revealed condition-specific microbial signatures: IPVB cases were characterized by dominance of Bacilli and Alphaproteobacteria, and PID cases showed enrichment of Actinomycetes and Alphaproteobacteria. In contrast, Infertility cases were distinguished by elevated Gammaproteobacteria. Collectively, pointing up the presence of systematic, condition-specific dysbiosis in the cervicovaginal microbiome, highlighting how distinct clinical presentations are associated with unique microbial shifts (Figure 4).

**Figure 4.**
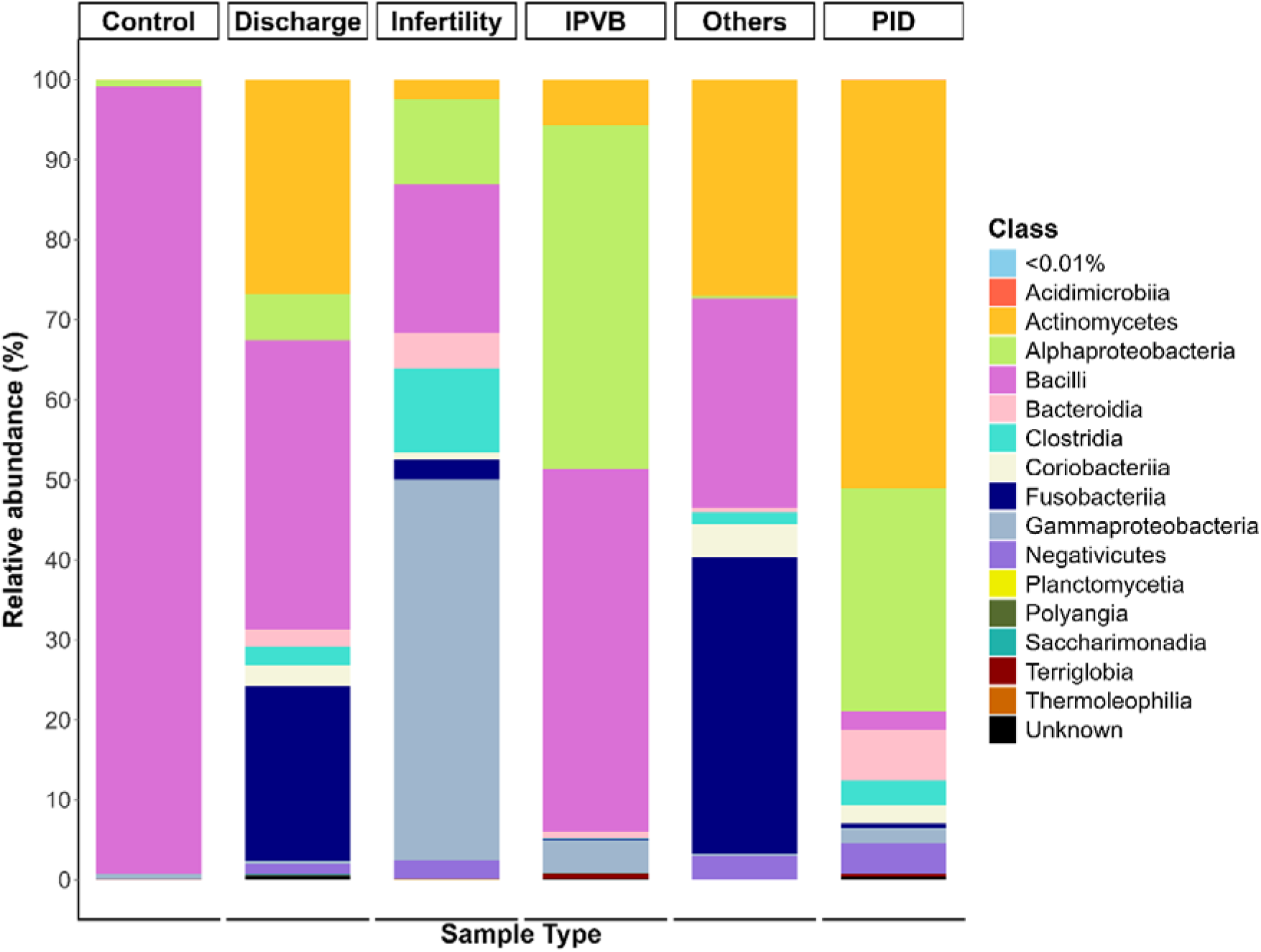
Relative abundance of cervicovaginal bacterial classes across pathological states.

#### 3.3.2 Family- and Genus-Level profile

At the family level, the cervicovaginal microbiome of healthy Controls was overwhelmingly dominated by Lactobacillaceae (98.38%), reflecting a stable, low-diversity community characteristic of protective *Lactobacillus*-driven ecosystems. In contrast, Pathological samples exhibited a sharp decline in Lactobacillaceae abundance (24.38%), accompanied by increased representation of Bifidobacteriaceae, Leptotrichiaceae, and Burkholderiaceae, indicating a transition toward a dysbiotic state. Subgroup-specific patterns were evident: Infertility cases were dominated by Burkholderiaceae (47.33%), PID cases showed elevated Bifidobacteriaceae (47.88%), IPVB cases were co-dominated by Bacillaceae and Caulobacteraceae, while Discharge and Other cases displayed mixed profiles enriched in diverse anaerobic taxa. These findings show condition-specific microbial signatures and highlight the ecological instability associated with pathological states.

Differential abundance analysis confirmed these compositional shifts. Lactobacillaceae was significantly enriched in Controls (log□FC = 3.35, padj = 0.0023), whereas Bifidobacteriaceae, Burkholderiaceae, Atopobiaceae, and Sphingomonadaceae were significantly enriched in Pathological groups (padj < 0.01). Core taxa analysis further revealed that Lactobacillaceae, Caulobacteraceae, and Xanthobacteraceae were consistently shared across groups, whereas several families, including Bifidobacteriaceae, Burkholderiaceae, and Sphingomonadaceae, were unique to Pathological samples. Visualization of these families in boxplots (Figure 5) and heatmaps highlighted both the depletion of protective taxa and the enrichment of disease-associated families, reinforcing the systematic shift toward a dysbiotic cervicovaginal microbiome.

**Figure 5.**
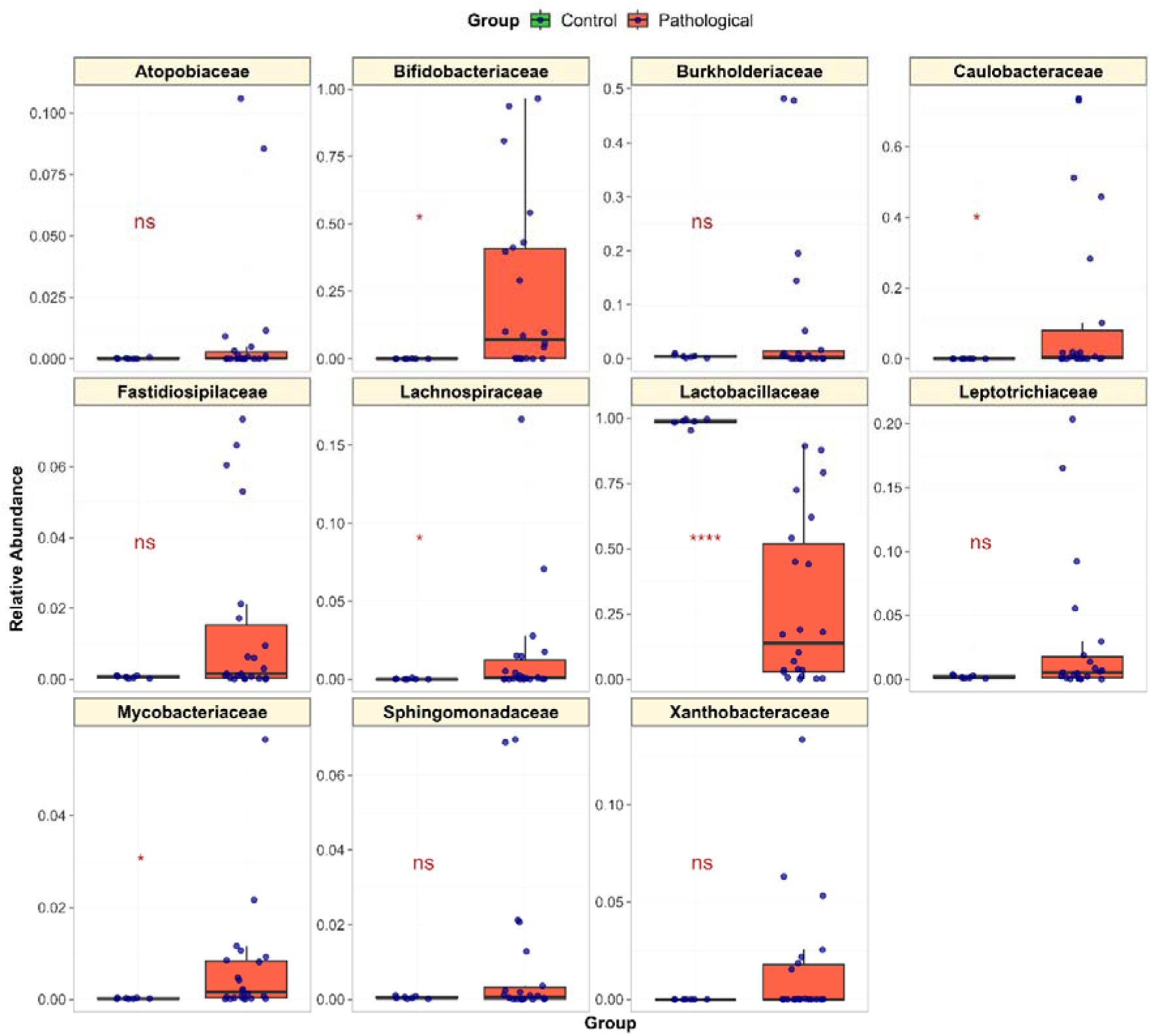
Relative abundance of core cervicovaginal microbial families in Control and Pathological groups.

At the genus level, the cervicovaginal microbiome of healthy Controls was dominated by Lactobacillus (98.2%), with only trace abundances of *Acinetobacter*, *Limosilactobacillus*, and *Mesorhizobium*. In contrast, Pathological samples exhibited a marked reduction in *Lactobacillus* (28.0%) and a substantial increase in potentially dysbiosis-associated genera, including *Bifidobacterium* (23.4%), *Achromobacter* (12.9%), and *Sneathia* (7.5%). These findings highlight a pronounced ecological shift from a *Lactobacillus*-dominated protective microbiome toward a more diverse and heterogeneous community in the disease state. Distinct genus-level signatures were evident across clinical subgroups. IPVB cases retained relatively high *Lactobacillus* abundance (58.2%), whereas PID cases showed a dramatic reduction (2.1%), with *Bifidobacterium* predominating (48.7%). Infertility samples were characterized by a striking enrichment of *Achromobacter* (45.5%), whereas Discharge and Other pathological groups displayed mixed profiles, with co-dominance of *Bifidobacterium*, *Sneathia*, and *Prevotella*. Together, these outcomes show condition-specific microbial dysbiosis, with distinct pathogenic or opportunistic genera driving divergence from the protective *Lactobacillus*-dominated state (Figure 6).

**Figure 6.**
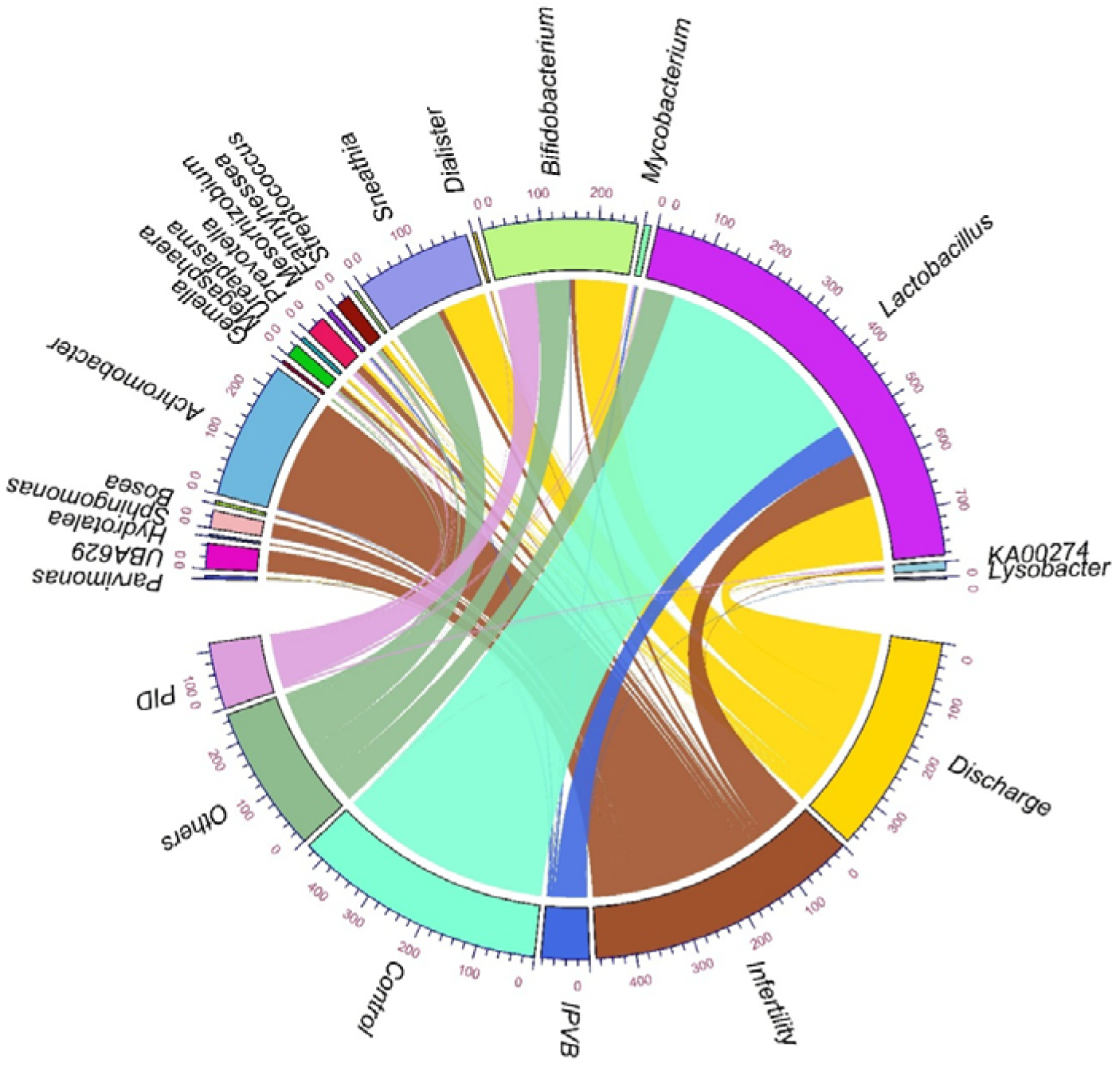
Genus-level distribution of cervicovaginal microbiota across clinical conditions.

#### 3.3.3 Species-level profiles

Species-level analysis further highlighted the divergence between healthy and pathological states. Across all samples, *Lactobacillus iners* (23.15%) emerged as the most abundant species, followed by *Bifidobacterium spp.* and *Sneathia sanguinegens.* Control samples were dominated by *L. iners* (74.07%) and other *Lactobacillus spp.* (23.83%), consistent with a protective, low-diversity microbial environment characteristic of cervicovaginal health. In contrast, Pathological samples showed a sharp reduction in *L. iners* (12.97%) and increased representation of anaerobic and opportunistic species. Remarkably, Infertility cases showed minimal *Lactobacillus* (9.66%) and were dominated by *Achromobacter spp.* strengthening the distinct dysbiosis signature observed at higher taxonomic levels, showing the progressive ecological shift from a *Lactobacillus*-dominated protective microbiome toward a heterogeneous community enriched in opportunistic taxa, particularly in infertility-associated dysbiosis.

### 3.4 Co-occurrence Network Analysis Reveals Dysbiosis-Associated Taxa

Genus-level correlation network analysis (Pearson correlation with FDR correction) uncovered distinct co-occurrence modules structured by clinical context. In healthy Controls, a tightly interconnected cluster was observed among *Mesorhizobium, Mycobacterium, Lysobacter, Ralstonia*, and *Reyranella*, with extremely strong positive correlations (r = 0.93-0.99, FDR < 10^-^¹²). This network reflects coordinated abundance patterns within control-dominant communities, consistent with ecological stability. By contrast, *Lactobacillus* exhibited consistent negative correlations with multiple dysbiosis-associated genera, including *Prevotella, Megasphaera, Parvimonas, Dialister,* and *KA00274* (r ≈ −0.33 to −0.41). However, these inverse associations did not remain statistically significant after FDR correction, suggesting compositional competition rather than robust ecological coupling.

In Pathological samples, a second highly significant module emerged among dysbiosis-associated genera. Strong positive correlations were detected between *Megasphaera, Prevotella, Parvimonas, Dialister*, and *KA00274* (r = 0.71-0.98, FDR < 10_-_□), forming a tightly co-occurring anaerobic consortium characteristic of disease-associated microbiota. Within infertility cases, a distinct cluster was observed among *Achromobacter, Sphingomonas, Bosea*, and *Hydrotalea* (r = 0.74-0.85, FDR < 10^-^□), reinforcing the infertility-specific dysbiosis signature. Additionally, *Streptococcus* showed a significant positive association with *Ureaplasma* (r = 0.65, FDR = 0.001), suggesting coordinated expansion in discharge-associated conditions. Collectively, these findings demonstrate modular organization of the cervicovaginal microbiome, with discrete, condition-specific co-occurrence networks underlying clinical stratification (Figure 7).

**Figure 7.**
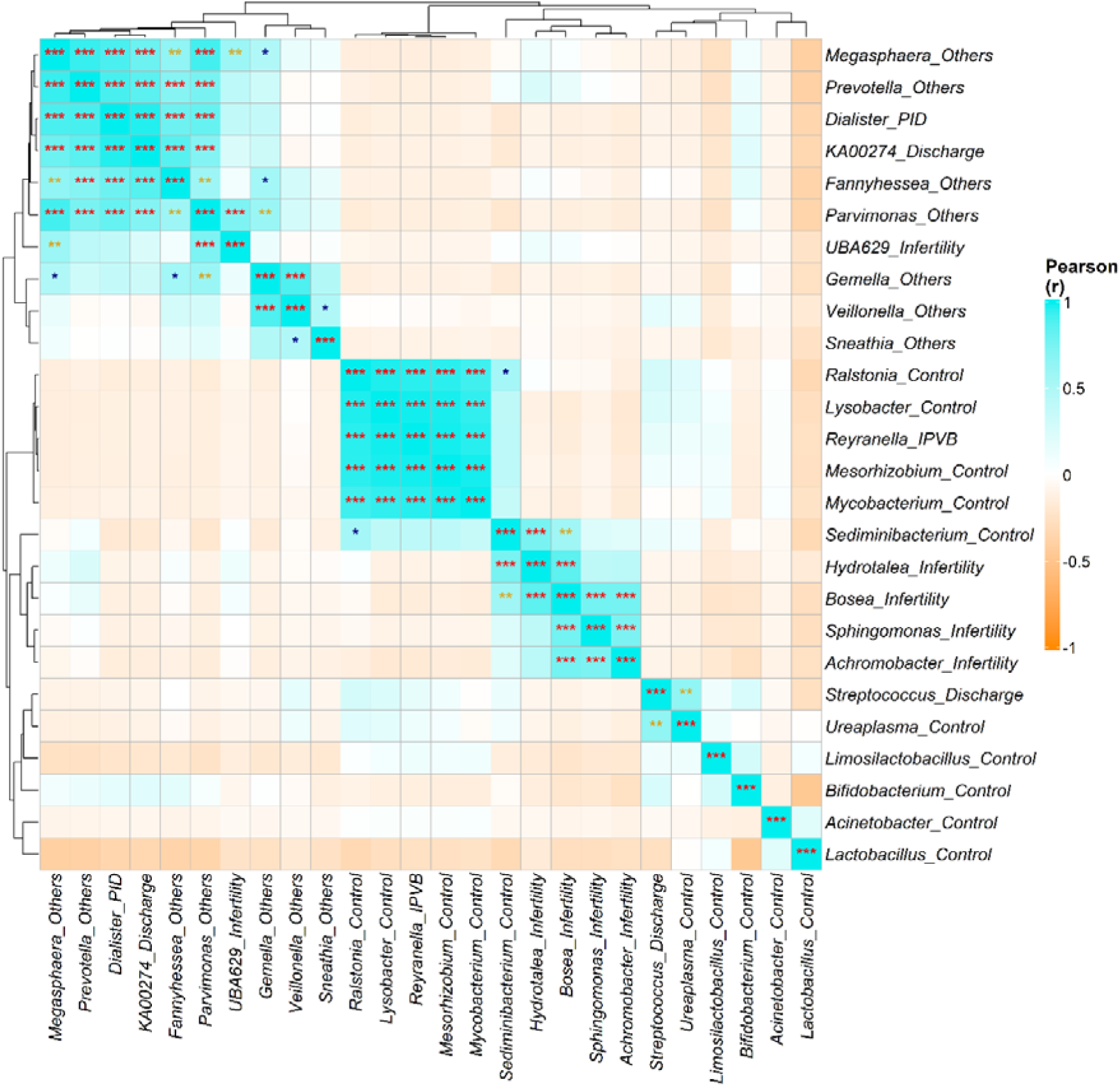
Genus-level co-occurrence networks across clinical types.

### 3.5 Functional Potential of the Cervicovaginal Microbiome

To investigate the predicted functional landscape of the cervicovaginal microbiome in HPV-negative women, PICRUSt2 was used to infer metabolic pathways from 16S rRNA gene profiles. Across all samples, a conserved core functional backbone was observed, dominated by carbohydrate metabolism, nucleotide biosynthesis, amino acid synthesis, and energy production. Pathways such as glycolysis (GLYCOLYSIS), pyruvate fermentation to lactate (FERMENTATION-PWY), and nucleotide biosynthesis modules were consistently abundant, reflecting the fundamental metabolic requirements of cervicovaginal microbial communities. This shared functional architecture indicates that, despite taxonomic variability, key metabolic processes remain preserved across health and disease states.

Differential pathway analysis using DESeq2 identified disease-specific functional enrichments (FDR < 0.05). Compared with Controls, Infertility samples showed significant enrichment of RUMP-PWY (log□FC = 5.37, padj = 0.003) and PWY-1861 (log□FC = 4.91, padj = 0.005). The shift was more pronounced in PID, where RUMP-PWY (log□FC = 8.16, padj = 3.16 × 10□□), PWY-1861 (log□FC = 7.61, padj = 4.60 × 10□□), and PWY-7013 (log□FC = 6.62, padj = 0.001) were strongly enriched, indicating intensified metabolic remodeling in inflammatory conditions. Discharge samples displayed the broadest functional expansion, with enrichment of central carbon metabolism and biosynthetic pathways, including GLYCOLYSIS (padj = 0.020), RIBOSYN2-PWY (padj = 0.011), COA-PWY-1 (padj = 0.016), and multiple carbohydrate and cell wall biosynthesis modules such as PEPTIDOGLYCANSYN-PWY (padj = 0.049), UDPNAGSYN-PWY (padj = 0.048), and PWY-6385/6386/6387 (padj < 0.05). Additional enrichment of LACTOSECAT-PWY, PHOSLIPSYN-PWY, POLYISOPRENSYN-PWY, and DAPLYSINESYN-PWY further supported enhanced carbohydrate utilization, membrane biosynthesis, and cell proliferation in discharge-associated microbiomes. Collectively, these findings indicate a stepwise intensification of metabolic and biosynthetic capacity from Infertility to PID and Discharge states, reflecting increased microbial growth, stress adaptation, and inflammatory engagement, while core metabolic functions remain partially conserved (Figure 8).

**Figure 8.**
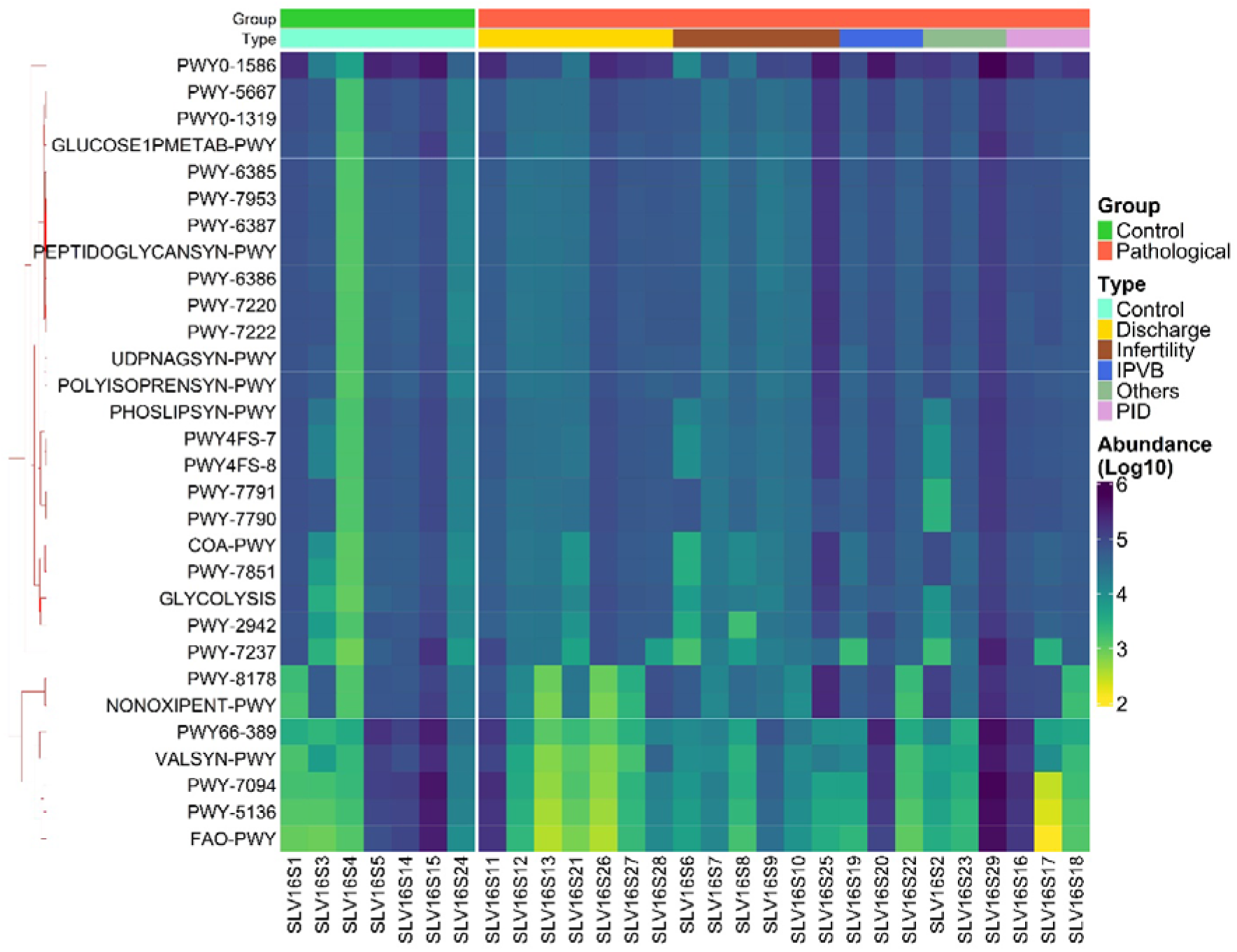
Predicted functional potential of the cervicovaginal microbiome in HPV-negative women.

### 3.6 Shotgun Metagenomics Confirms *Achromobacter* Dominance in Infertility

#### 3.6.1 Viral and Bacterial Composition

Shotgun metagenomic sequencing of three infertility-associated samples provided high-resolution insights into the cervicovaginal ecosystem. The bacterial community was overwhelmingly dominated by Proteobacteria (96.4%), with Firmicutes and Actinobacteria present at low levels. Viral taxa accounted for 4.40% of total abundance, primarily human endogenous retrovirus K113, followed by orthobunyaviruses and bacteriophages. These findings suggest potential viral-bacterial interactions within the dysbiotic cervicovaginal niche (Figure 9).

**Figure 9.**
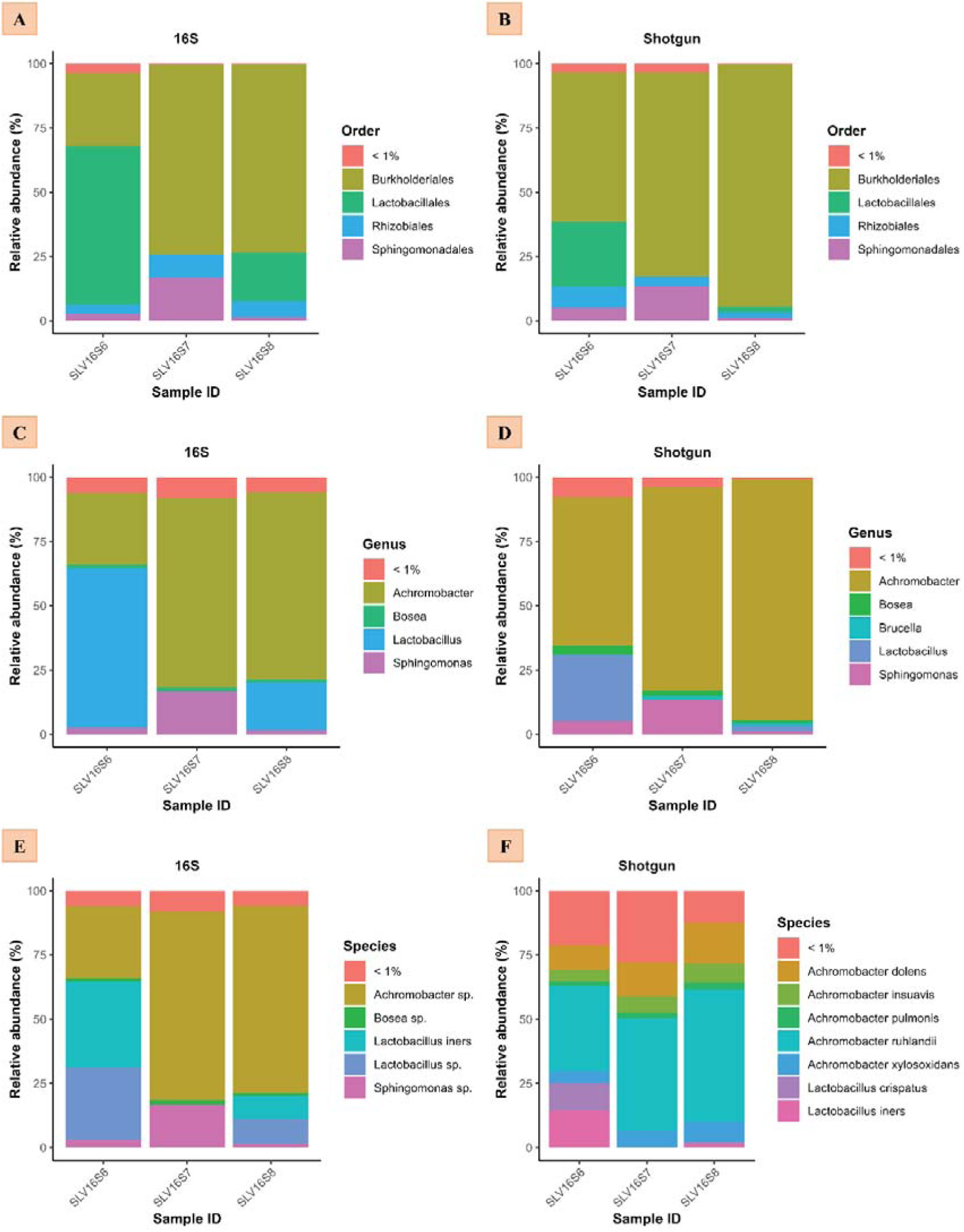
Shotgun versus amplicon-based taxonomic profiles in infertility samples.

#### 3.6.2 High-Resolution Taxonomic Profiling

At the order level, Burkholderiales dominated across all samples (58–94%). Within this order, *Achromobacter* was the most abundant genus (84.9%), with species-level resolution identifying *A. ruhlandii*, *A. dolens,* and *A. xylosoxidans* as the principal taxa. In contrast, *Lactobacillus* was considerably depleted, particularly in one sample where it accounted for only 2.21% of total abundance. These results strengthen the severe loss of protective *Lactobacillus* species and the emergence of *Achromobacter* as a dominant taxon in infertility-associated dysbiosis.

#### 3.6.3 Concordance with Amplicon-Based Profiling

Shotgun metagenomics validated the 16S rRNA gene sequencing results, confirming *Achromobacter* as the dominant taxon in infertility samples and providing species-level resolution absent from amplicon data. This cross-platform concordance strengthens the evidence for *Achromobacter* as a potential microbial biomarker of infertility. It highlights the value of integrating amplicon and shotgun approaches for comprehensive cervicovaginal microbiome profiling.

Taken together, the shotgun metagenomic data provide high-resolution confirmation of the *Achromobacter*-dominated dysbiosis observed in infertility cases. The concordance between amplicon-based and shotgun approaches strengthens the robustness of this finding, while species-level resolution highlights the predominance of *A. ruhlandii, A. dolens,* and *A. xylosoxidans*. The severe depletion of protective *Lactobacillus* species, coupled with the expansion of opportunistic *Achromobacter*, underscores profound ecological disruption in the cervicovaginal niche. Importantly, the cross-platform validation positions *Achromobacter* as a potential microbial biomarker of infertility, suggesting translational relevance for diagnostics and targeted interventions. These results highlight the value of integrating amplicon and shotgun sequencing to capture both community-level shifts and species-specific signatures, thereby advancing our understanding of cervicovaginal dysbiosis in reproductive health.

Collectively, our findings reveal a progressive ecological and functional disruption of the cervicovaginal microbiome in HPV-negative women with pathological conditions. Diversity analyses demonstrated significantly higher alpha diversity and greater beta-diversity dispersion in pathological groups, reflecting destabilization of the protective *Lactobacillus*-dominated community. Taxonomic profiling across multiple levels consistently highlighted a shift from Bacillota/Bacilli dominance in Controls toward enrichment of Actinomycetota, Fusobacteriota, Gammaproteobacteria, and Bifidobacteriaceae in disease states, with condition-specific signatures such as Achromobacter in infertility and Bifidobacterium in PID. Functional inference revealed a conserved metabolic backbone across all samples but identified disease-specific enrichment of biosynthetic and stress-adaptation pathways, particularly in PID and discharge-associated microbiomes. Co-occurrence network analysis further demonstrated modular organization, with tightly interconnected control-associated clusters contrasting with anaerobic consortia and *Achromobacter*-centered modules in pathological states. Finally, shotgun metagenomics provided species-level resolution, confirming the dominance of *Achromobacter* spp. (*A. ruhlandii, A. dolens, A. xylosoxidans*) in infertility samples and validating amplicon-based findings. Together, these results underscore a consistent pattern of taxonomic dysbiosis, functional remodeling, and condition-specific microbial interactions, positioning Achromobacter as a potential biomarker of infertility and highlighting the translational value of integrated multi-omics approaches in cervicovaginal microbiome research.

## 4. Discussion

This study provides an integrated characterization of cervicovaginal microbiome dynamics in HPV-negative women with diverse gynecological conditions, combining 16S rRNA gene sequencing with shotgun metagenomics to achieve both breadth and taxonomic resolution. While cervicovaginal dysbiosis has been extensively investigated in HPV-positive women, microbial shifts in HPV-negative individuals remain comparatively underexplored, particularly in low-resource settings. Our findings reveal consistent patterns of increased microbial diversity, depletion of protective *Lactobacillus* species, and enrichment of anaerobic and opportunistic taxa across pathological states. Notably, infertility cases exhibited a striking dominance of *Achromobacter spp.*, validated across sequencing platforms, positioning this genus as a potential biomarker of reproductive dysfunction. These results underscore the ecological instability of the cervicovaginal niche in disease conditions and highlight the translational importance of microbiome-informed diagnostics and therapeutic strategies aimed at restoring reproductive health.

### 4.1 Ecological Interpretation of Cervicovaginal Dysbiosis

Our results demonstrate that cervicovaginal dysbiosis in HPV-negative women is characterized by increased microbial diversity, loss of *Lactobacillus* dominance, and enrichment of anaerobic and opportunistic taxa. The rise in alpha diversity and dispersion in pathological groups reflects ecological destabilization, consistent with previous reports linking microbial heterogeneity to mucosal inflammation and reproductive dysfunction. Condition-specific signatures such as *Achromobacter* in infertility, *Bifidobacterium* in PID, and mixed anaerobic consortia in discharge cases-underscore that dysbiosis is not uniform but stratified by clinical presentation. Co-occurrence network analysis further revealed modular organization, with tightly interconnected clusters in healthy controls contrasting with anaerobic consortia in disease states. These findings suggest that cervicovaginal dysbiosis involves both taxonomic shifts and altered microbial interactions, reinforcing the concept of ecological imbalance as a driver of gynecological pathology.

### 4.2 Clinical Implications and Biomarker Potential

The cross-platform validation of *Achromobacter* dominance in infertility cases positions this genus as a candidate biomarker for reproductive dysfunction. Species-level resolution provided by shotgun metagenomics identifying *A. ruhlandii, A. dolens,* and *A. xylosoxidans* adds translational value by pinpointing taxa that may be clinically relevant. The depletion of protective *Lactobacillus* species, coupled with the expansion of opportunistic *Achromobacter*, highlights a profound ecological disruption that may compromise mucosal immunity and reproductive outcomes. Functional inference further revealed enrichment of biosynthetic and stress-adaptation pathways in pathological states, suggesting enhanced microbial growth and inflammatory engagement. Together, these findings support the potential utility of microbiome-informed diagnostics and interventions, particularly in the management of infertility. However, the limited sample size warrants cautious interpretation, and larger, longitudinal studies are needed to validate *Achromobacter* as a reliable biomarker and to explore its mechanistic role in reproductive health.

### 4.3 Limitations and Future Directions

While this study provides novel insights into the dynamics of the cervicovaginal microbiome in HPV-negative women, several limitations must be acknowledged. First, the relatively small sample size particularly for shotgun metagenomics (n = 3 infertility cases) limits statistical power and generalizability. Findings should therefore be interpreted as preliminary, requiring validation in larger, multi-centre cohorts. Second, the cross-sectional design precludes causal inference; observed associations between microbial shifts and clinical conditions may reflect correlation rather than direct mechanistic effects. Third, functional predictions derived from 16S rRNA gene data (PICRUSt2) provide valuable hypotheses but remain inferential, underscoring the need for direct metatranscriptomic or metabolomic validation. Finally, host factors such as hormonal status, sexual behavior, and antibiotic exposure were not fully integrated into the analysis, and these variables may significantly shape cervicovaginal ecology.

Future research should prioritize longitudinal studies to capture the temporal dynamics of dysbiosis, particularly in relation to infertility and inflammatory conditions. Expanding shotgun metagenomic sequencing across diverse clinical subgroups will enable deeper resolution of microbial taxa and functional pathways, while integration with metabolomic and immunological profiling could clarify host-microbe interactions. Importantly, the consistent dominance of *Achromobacter* in infertility cases warrants targeted investigation into its pathogenic potential, ecological role, and suitability as a biomarker for reproductive dysfunction. Such studies could inform the development of microbiome-based diagnostics and therapeutic interventions aimed at restoring cervicovaginal health.

The cervicovaginal microbiome plays a critical role in reproductive health, yet its dynamics in HPV-negative women remain poorly understood. By integrating amplicon and shotgun metagenomics, this study reveals profound ecological disruption in pathological conditions, marked by loss of protective *Lactobacillus* and emergence of *Achromobacter* as a dominant taxon in infertility. These findings not only expand our understanding of cervicovaginal dysbiosis but also identify *Achromobacter* as a candidate biomarker with translational relevance for infertility diagnostics. The study highlights the importance of multi-omics approaches in uncovering condition-specific microbial signatures and sets the stage for future work on microbiome-targeted interventions to improve reproductive outcomes.

## 5. Conclusion

This study provides the first integrated characterization of cervicovaginal microbiome dynamics in HPV-negative women across diverse gynaecological conditions, combining amplicon-based 16S rRNA sequencing with shotgun metagenomics. Our results reveal a consistent pattern of increased microbial diversity, depletion of protective *Lactobacillus* species, and enrichment of anaerobic and opportunistic taxa in pathological states. Particularly, infertility cases were marked by striking dominance of *Achromobacter spp.*, validated across sequencing platforms, positioning this genus as a potential biomarker of reproductive dysfunction. Functional inference highlighted both conserved metabolic backbones and disease-specific enrichment of biosynthetic and stress-adaptation pathways, reflecting progressive ecological remodelling. Co-occurrence network analysis further demonstrated condition-specific microbial interactions, underscoring the modular organization of dysbiosis. Collectively, these findings advance our understanding of cervicovaginal ecology in HPV-negative women and emphasize the translational potential of microbiome-informed diagnostics and therapeutic strategies in reproductive health.

## 6. Data availability

The 16S and shotgun sequence data are available in the NCBI database under the BioProject.

